# VascFlexMap: Microvascular Ultrasound Imaging at Low Frame Rates Using Sparse Data and a Transformer-Decoder Network

**DOI:** 10.64898/2026.02.27.708398

**Authors:** Ruchika Dhawan, Mihir Agarwal, Shreyans Jain, Himanshu Shekhar

## Abstract

**Objective:** Super-resolution ultrasound (SR-US) reveals microvascular structures with exquisite resolution, but clinical translation remains limited by the need for ultrafast frame rates, massive data volumes, and long reconstruction times. This work proposes a deep learning framework that reconstructs microvascular maps from low-frame-rate enhanced ultrasound sequences, bypassing explicit microbubble localization and tracking.

**Methods:** A transformer-decoder network with learned linear projections was designed to model spatiotemporal dependencies across sparse contrast-enhanced ultrasound sequences and reconstruct vessel probability maps, refined via a post-processing enhancement stage. Single-head self-attention captures temporal correlations under challenging conditions including overlapping microbubbles and low signal-to-noise ratios. Binary cross-entropy loss guided training to preserve vascular topology across synthetic and *in vivo* datasets. *In vivo* rat brain bolus data from the PALA challenge was used to evaluate this approach under up to 500 − fold data reduction (341 frames at 2 FPS vs. 170400 frames at 1000 FPS in standard ULM).

**Results:** Despite aggressive undersampling, the proposed pipeline recovered coherent microvascular architecture where conventional ULM pipelines applied to the same sparse data failed to produce continuous vascular networks. Major branches and higher-order microvessels remained visible with apparent vessel widths broadened by approximately three-fold relative to reference SR-US. End-to-end reconstruction completed in 28–133 seconds on an NVIDIA H100 GPU depending on the number of frames employed.

**Conclusion:** The reported approach preserved vascular topology with fast reconstruction and low data overhead, albeit at lower resolution. The substantial reduction in frames and computation time highlights the translational potential of this SR-US-inspired microvascular imaging approach.

## 1 Introduction

Microvasculature imaging is under development for the assessment of early pathological changes in cancer, stroke, diabetes, and inflammation, where microvascular alterations precede macroscopic structural damage [1–3]. Conventional power Doppler ultrasound operates at low frame rates (typically *<*50 Hz) and offers poor spatial resolution for vessels below approximately 0.5 - 1 mm diameter due to limited signal-to-noise ratio, suppression of slow flows (*<*1-5 mm/s) by wall filters, and high susceptibility to tissue motion artifacts arising from breathing or cardiac pulsation [4–6]. Although power Doppler remains clinically ubiquitous for mapping larger vessels (*>*500 *µ*m), it fails to resolve the capillary beds that account for more than 90% of tissue perfusion, thus limiting its sensitivity to early pathological changes in oncologic and inflammatory disease processes [7].

Ultrafast Doppler achieves higher microvascular sensitivity through compounded imaging at elevated frame rates (500-1000 Hz) that suppress clutter via temporal averaging [8,9]. This yields spatial resolution that is intermediate between conventional Doppler and super-resolution ultrasound (SR-US) but requires specialized high-frame-rate hardware and cannot typically resolve sub-100 *µ*m capillaries due to persistent diffraction limits.

Contrast enhanced ultrasound (CEUS) leverages encapsulated microbubbles as nonlinear acoustic tracers [10] to enhance visualization of blood vessels [11–16]. These gas-filled microbubbles [17], typically smaller than 5 *µ*m in diameter, exhibit strong nonlinear scattering when insonified, improving signal-to-noise ratio and contrast [18,19]. The main limitation of this approach is its limited resolution. Super-resolution ultrasound (SR-US), particularly ultrasound localization microscopy (ULM), attains true microvessel-scale resolution (*<*50 *µ*m) by repeatedly imaging, localizing via sub-pixel centroid fitting, and tracking individual microbubbles across thousands of frames (10,000-100,000 per volume) acquired at ultrafast rates [20–22]. However, this approach imposes massive data storage requirements (several GBs per exam), exorbitant computational loads for sequential denoising, detection, association, tracking, and motion compensation (typically hours of post-processing), and dependency on research-grade systems with ultrafast capabilities incompatible with clinical scanners limited to modest frame rates (~25-80 Hz), and data bandwidth [23–25].

These fundamental bottlenecks, including requirement for ultrafast acquisition, data/storage overhead, and offline processing, severely hinder clinical translation of SR-US [26].

Deep learning approaches have accelerated SR-US by learning direct end-to-end mappings from CEUS data to vascular probability maps, bypassing computationally expensive localization pipelines [27, 28]. While convolutional neural networks (CNNs) like Deep-ULM improve localization speed, they typically assume relatively dense temporal sampling and struggle to capture long-range spatiotemporal dependencies critical for sparse, low-frame-rate inputs [29, 30]. Transformer architectures excel at global context modeling through self-attention, and hybrid encoder-decoder transformer networks have demonstrated superior edge-preserving segmentation under data scarcity [31–34].

Importantly, the application of microvascular imaging in the clinical setting may not need exact micron-level capillary recovery, but rather the rapid depiction of vascular topology. In routine clinical workflows, actionable insight within minutes is important, even if tade offs need to be considered with respect to spatial resolution. Accordingly, lower-resolution yet rapidly reconstructed microvascular maps may still be acceptable for clinical decision-making and could also be used to identify regions warranting targeted high-fidelity SR-US acquisition.

This work proposes **VascFlex Map**, a transformer-decoder network with learned linear projections for robust microvascular reconstruction from low-frame-rate (~2-50 Hz) sparse CEUS sequences. Evaluations on *in vivo* rat brain bolus data from the PALA challenge demonstrate preservation of major vascular branches and higher-order microvessels down to 500× data reduction (341 frames at 2 FPS vs. 170400 frames ULM), achieving −3 dB FWHM of 110.88 ± 15.23 *µ*m versus ULM 34.91 ± 30.46 *µ*m. End-to-end inference completed in 28–133 s on NVIDIA H100 GPUs, advancing sparsity-tolerant, hardware-agnostic SR-US toward practical clinical deployment for applications such as stroke triage, tumor margin assessment, and cancer imaging.

## 2 Methods

### 2.1 Problem Formulation

Super-resolution ultrasound localization microscopy requires ultrafast frame rates (over 1000 FPS) to capture individual microbubble trajectories. This work addresses the inverse problem: given only a sparse temporal sequence of *N* frames acquired at clinically feasible frame rates (2–50 FPS), reconstruct a vessel probability map that preserves essential microvascular topology. Rather than explicitly detecting and tracking individual microbubbles, the problem is formulated as a learned mapping from sparse temporal observations to dense vascular structure, leveraging deep learning to implicitly model the spatiotemporal dynamics of contrast agent flow. Fig. 1 illustrates the end-to-end processing pipeline.

**Figure 1.**
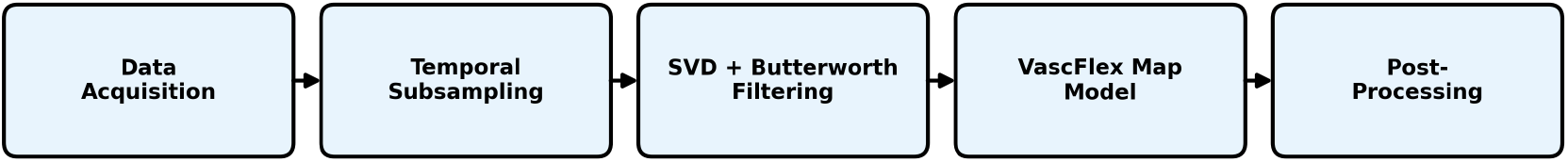
End-to-end processing pipeline. B mode images acquired at 1000 Hz undergo temporal subsampling to clinical frame rates (2–50 Hz), followed by SVD-based clutter suppression and Butterworth bandpass filtering. The VascFlex Map model processes filtered sequences to generate vessel probability maps, which were refined through post-processing and upscaled to 8192×8192 resolution.

### 2.2 Dataset Acquisition

The publicly available rat brain bolus imaging dataset from the PALA challenge [24] was used to evaluate the proposed framework under physiologically relevant conditions. This dataset contains high-frequency (15 MHz) contrast-enhanced ultrasound acquisitions of rat brain microvasculature obtained following intravenous bolus injection of Sonovue microbubbles. Data were acquired using a linear array probe (128 elements, *λ*-pitch) with compounded plane wave imaging (3–5 tilted angles) at frame rates up to 1,000 Hz over a 7×14.9 mm^2^ field of view. Each acquisition contains 213 blocks of 800 frames, providing dense temporal sampling for ground-truth super-resolution reconstructions via conventional ultrasound localization microscopy (ULM).

### 2.3 Data Preparation and Preprocessing

All *in vivo* rat brain bolus datasets underwent a multi-stage preprocessing pipeline prior to model input. First, temporal subsampling was applied to the raw IQ (in-phase/quadrature) data to simulate clinical frame rate constraints. Every *N* th frame was systematically selected from each 800-frame acquisition block to achieve target effective frame rates of 50 Hz (*N* =20), 25 Hz (*N* =40), 15 Hz (*N* =66), 5 Hz (*N* =200), 3 Hz (*N* =333), and 2 Hz (*N* =500), representing up to 500-fold data reduction from the original 1000 Hz ultrafast acquisition. Additionally, a last-*N* frames analysis was conducted to assess temporal sufficiency requirements, wherein only the final portion of each acquisition was retained at fixed effective frame rates, allowing systematic evaluation of how reconstruction quality varies with temporal coverage.

Following temporal subsampling, singular value decomposition (SVD)-based clutter suppression was applied to the spatiotemporally organized IQ data matrix. The SVD filter decomposes the data into orthogonal singular components, where low-rank components correspond to stationary or slowly moving tissue clutter and higher-rank components capture the sparse, rapidly varying microbubble signals. By discarding singular values below a threshold (retaining indices 2 through *N*_frames_), the filter effectively removes tissue background while preserving microbubble signatures essential for vascular reconstruction.

A second-order Butterworth bandpass filter was subsequently applied along the temporal dimension to further suppress residual clutter and isolate microbubble dynamics within the physiologically relevant frequency range. The bandpass cutoff frequencies were scaled according to the effective frame rate after subsampling, with the filter applied only when the upper cutoff remained below the Nyquist frequency. This temporal filtering attenuates both low-frequency tissue motion artifacts and high-frequency noise outside the expected microbubble velocity spectrum.

Envelope-detected images were then resized to 256×256 resolution for input to the neural network pipeline. This cascaded preprocessing of subsampling, SVD filtering, and Butterworth band-pass filtering ensured optimal separation of microbubble signals from tissue background across all tested sparsity regimes.

### 2.4 Unconditional Learning Framework

A key innovation of the proposed approach is the unconditional learning formulation, wherein the network learns to reconstruct vascular structure without requiring actual ultrasound frames as input during inference. During training, synthetic input signals, specifically Gaussian random noise tensors of dimension (sequence length × 256 × 32 × 32), were paired with ground truth vascular maps derived from full-data ULM reconstructions. This unconditional formulation forced the network to learn robust vascular priors directly from the ground truth supervision rather than extracting features from noisy ultrasound data. The model effectively learned a generative mapping from random initialization to anatomically plausible vascular structure, conditioned only on the statistical patterns present in the training distribution. This approach aims to eliminate the sensitivity to acquisition-specific variations such as gain settings, tissue attenuation, and microbubble concentration fluctuations that can hamper reconstruction using conventional methods.

### 2.5 Network Architecture

The proposed VascFlex Map architecture consists of three main components: an input projection module, a temporal transformer encoder, and a transposed convolutional decoder. Fig. 2 illustrates the end-to-end pipeline.

**Figure 2.**
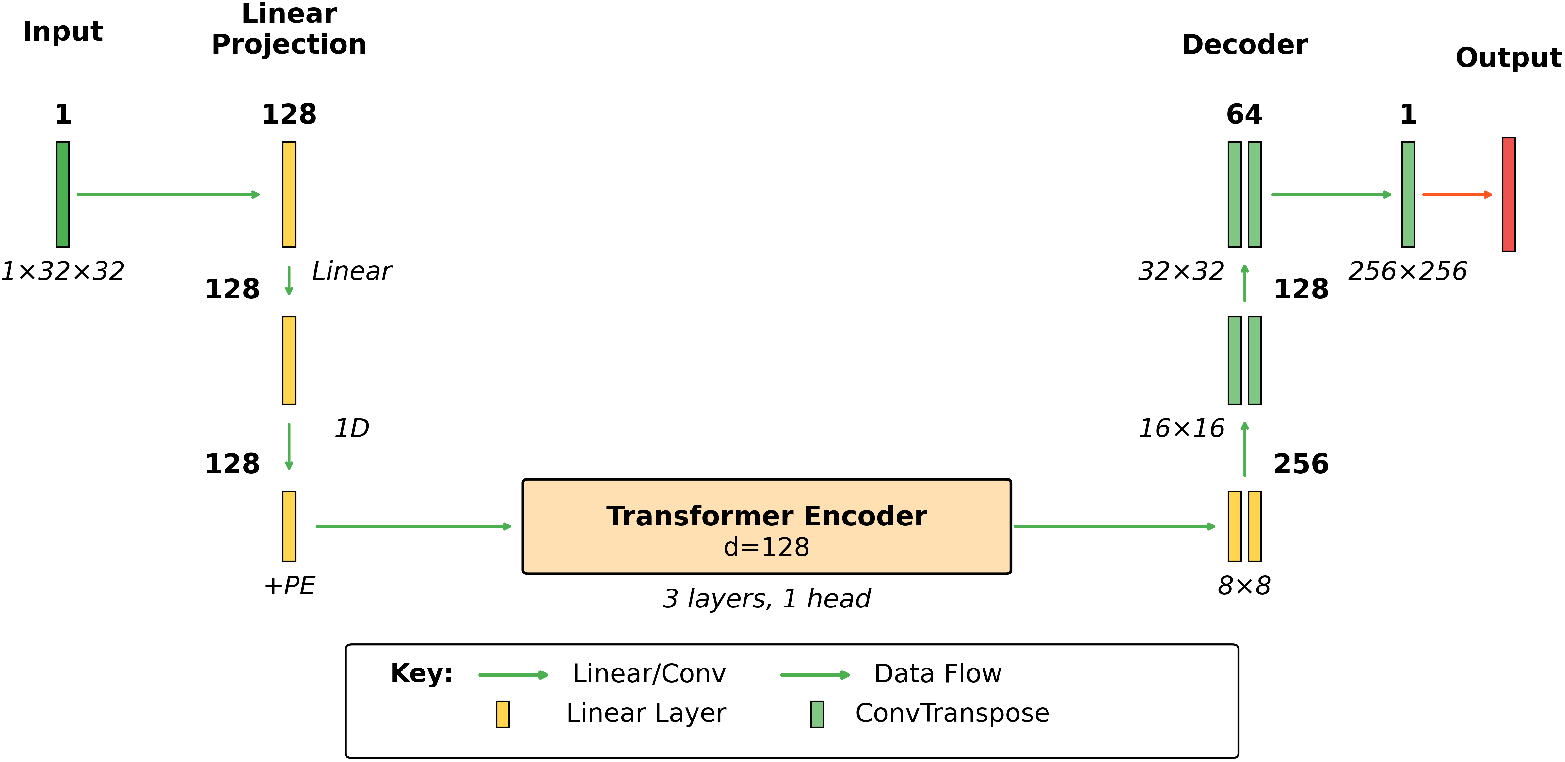
Proposed VascFlex Map architecture. Gaussian noise input of shape (sequence length × 256 × 32 × 32) was flattened and projected to 128 dimensions via a learned linear layer, processed by a transformer encoder with sinusoidal positional encoding (3 layers, 1 attention head), projected back to the original dimensions, and decoded through transposed convolutions (256→128→64→32→1 channels) to produce 256×256 vessel probability maps via sigmoid activation.

The input projection module flattened the synthetic input tensor of shape (sequence length × 256 × 32 × 32) to a vector of 262,144 dimensions and projected it to a lower-dimensional embedding space of *d*_model_ = 128 dimensions via a learned linear transformation. This compressed representation served as input to the temporal transformer encoder. The transformer encoder employed sinusoidal positional encoding to inject temporal order information into the sequence representation [31]. The positional encoding follows the standard formulation:

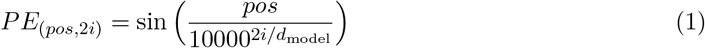

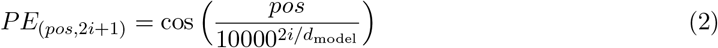

where *pos* denotes the temporal position and *i* indexes the embedding dimension. The encoded sequence was processed by three stacked transformer encoder layers, each containing single-head self-attention followed by position-wise feedforward networks. The self-attention mechanism enables the model to capture long-range temporal dependencies across the sparse frame sequence, learning which temporal positions contain complementary vascular information.

Following the transformer, a second linear projection mapped the 128-dimensional encoded representation back to 262,144 dimensions, which was then reshaped to (256 × 32 × 32) for decoding. The decoder consisted of three transposed convolution stages that progressively upsampled the feature maps from 32×32 to 256×256 resolution. Each stage comprised a transposed convolution with stride 2 for 2× upsampling, followed by a standard convolution with ReLU activation for feature refinement. The channel dimensions progress as 256→128→64→32 across the three stages. A final 1×1 convolution reduced the 32-channel output to a single-channel vessel probability map, and a sigmoid activation constrained the output to the range [0, 1], representing per-pixel vessel presence probability.

### 2.6 Loss Function

The network was trained using binary cross-entropy (BCE) loss, which measured the pixel-wise discrepancy between predicted vessel probability maps and ground truth binary segmentations. For a predicted probability map 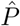 and ground truth binary mask *P*, the BCE loss is defined as:

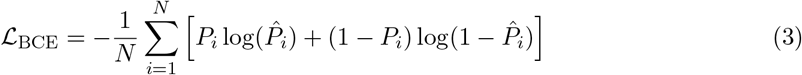

where *N* denotes the total number of pixels and *i* indexes individual pixel locations. This formulation penalizes confident incorrect predictions exponentially; a prediction of 0.01 for a true vessel pixel incurs substantially greater loss than a prediction of 0.4, thereby encouraging the network to output well-calibrated probability estimates. Binary cross-entropy is particularly well-suited for the vessel segmentation task as it treats each pixel as an independent binary classification problem, allowing the network to learn sharp vessel boundaries. The sigmoid activation in the final decoder layer ensured predictions remain in the valid probability range [0, 1], preventing numerical instability during loss computation [35].

### 2.7 Training Protocol

The network was trained using the Adam optimizer with a learning rate of 3 × 10^*−*4^, leveraging adaptive moment estimation to handle varying gradient magnitudes across the network’s linear and convolutional components [36]. Training proceeded with a batch size of 1 to accommodate memory requirements of processing full image sequences. The unconditional formulation enabled rapid convergence within 2 training epochs, as the network need not learn complex feature extraction from noisy ultrasound data but rather directly associates random initializations with target vascular patterns. Ground truth images were resized to 256×256 pixels and converted to single-channel grayscale format. The sampling rate parameter controls temporal sparsity by selecting every *N* th frame from the original acquisition, simulating the effect of reduced frame rate acquisition.

### 2.8 Post-Processing Pipeline

The raw network output underwent a multi-stage post-processing pipeline to enhance vessel visibility and prepare the final high-resolution reconstruction. First, Total Variation (TV) denoising with weight parameter 0.05 was applied using the Chambolle algorithm to suppress noise while preserving sharp vessel edges, to maintain vascular boundary definition [37].

Contrast-limited adaptive histogram equalization (CLAHE) was subsequently applied with a clip limit of 0.01 to enhance local contrast without amplifying noise in homogeneous background regions. This adaptive approach divides the image into tiles and equalizes each independently, preventing vessels in low-contrast regions from being obscured by dominant high-intensity structures elsewhere [38].

Unsharp masking with radius 1.0 and amount 1.2 was used to sharpen vessel edges by subtracting a blurred version of the image and adding the high-frequency residual back with amplification. This step recovered edge definition potentially softened by the denoising stage. Morphological erosion was performed with a 3×3 rectangular kernel to thin vessel structures by removing boundary pixels, partially compensating for the resolution broadening inherent in sparse-data reconstruction. Finally, the processed 256×256 output was upscaled by a factor of 32 to produce 8192×8192 pixel images suitable for detailed clinical inspection.

### 2.9 Quantitative Evaluation

Reconstruction fidelity was assessed using Full Width at Half Maximum (FWHM) resolution estimates. Intensity profiles were extracted across five manually selected vessel cross-sections in both reconstructed and reference images using identical spatial locations. Profiles were converted to decibel scale via 20 log_10_(*I/I*_max_), and FWHM was measured at the −3 dB contour. Wall-clock processing time was recorded for each subsampling condition on an NVIDIA H100 GPU to characterize computational efficiency.

### 2.10 Statistical Analysis

Statistical analysis was performed on the contrast values measured from five independent line profiles extracted from the raw and post-processed images. Normality of the paired differences was assessed using the Shapiro–Wilk test. If the normality assumption was satisfied, a two-sided paired *t*-test was used to compare contrast values before and after postprocessing. For non-normal data, the non-parametric Wilcoxon signed-rank test was applied. All statistical tests were two-sided with a significance level of *α* = 0.05. Data are reported as mean ± standard deviation.

## 3 Results

Fig. 3 illustrates the effect of the proposed post-processing pipeline on the model output for low-frame-rate acquisition data. The left panel shows the raw (pre-processed) probability map produced directly by the network from 25 FPS input (4260 frames), while the right panel shows the post-processed result after application of the enhancement pipeline. The post-processing significantly improves visual clarity by reducing background noise, enhancing local contrast, and sharpening vascular boundaries. Fine microvascular structures become more continuous and better defined, while spurious low-intensity artifacts are suppressed. Importantly, the processing preserves the underlying vascular topology while improving perceptual quality and interpretability. The mean contrast achieved at 25 FPS with the model before and after preprocessing were 0.3560 ± 0.2070 and 0.4680 ± 0.2439, respectively. However, the observed increase in contrast after preprocessing was not statistically significant (*p* = 0.4050; *p* ≥ 0.05) compared to the raw model output.

**Figure 3.**
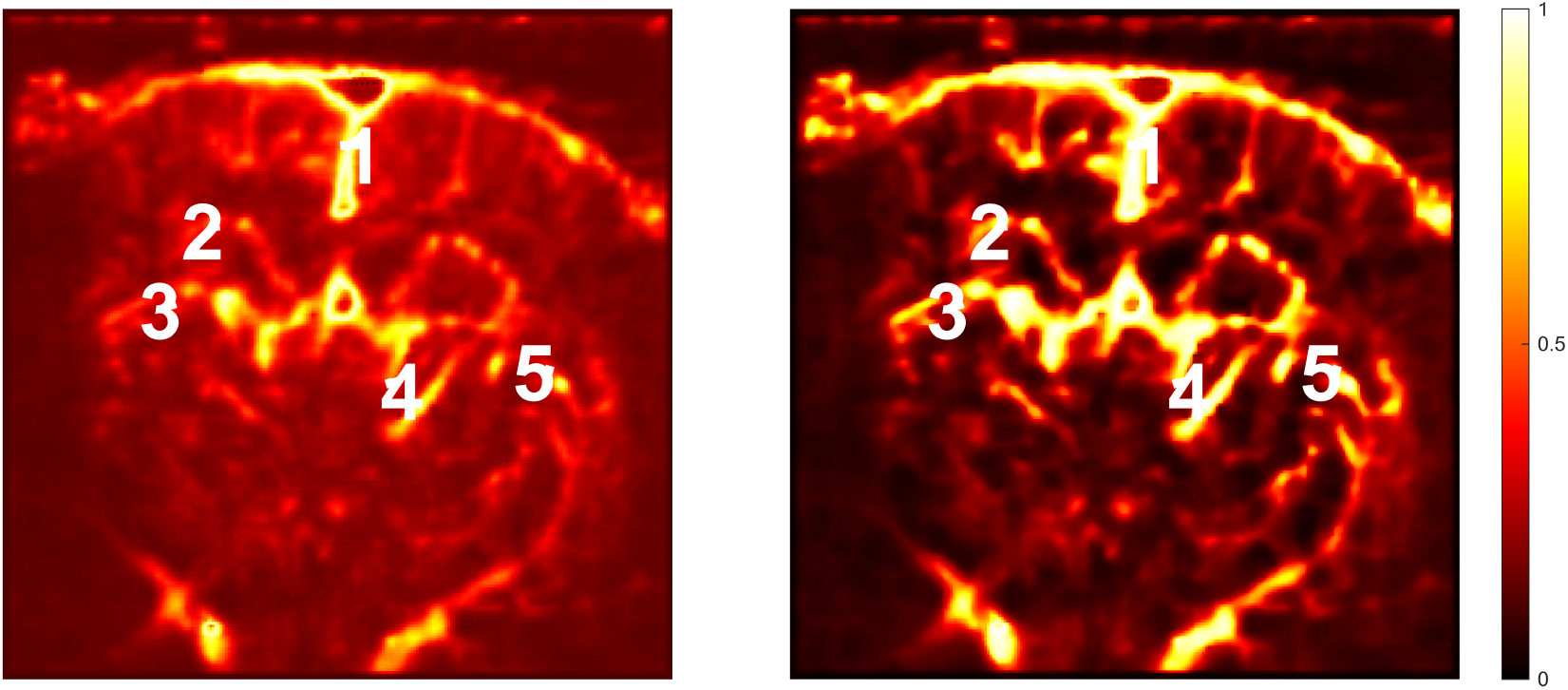
Post-processing effect on model output probability maps using low-frame-rate input. (Left) Raw network output from 25 FPS acquisition (4260 frames). (Right) Post-processed visualization after TV, CLAHE contrast enhancement, and unsharp masking.

Fig. 4 demonstrates post-processed reconstruction using the reported framework across highly sparse regimes using rat brain bolus data. Row 1 shows the conventional full-data ULM reconstruction (1000 FPS, 170400 frames). Rows 2–3 present VascFlex Map reconstructions at progressively lower frame rates: 50/25/15 FPS (4260–8520 frames) and 5/3/2 FPS (341–852 frames), representing 20–500× data reduction. Despite severe undersampling, vascular topology remains visible down to 2 FPS, with clear delineation of cortical branches and penetrating arterioles. Finer capillaries blur progressively (as expected), but major network architecture persists even at minimum viable inputs.

**Figure 4.**
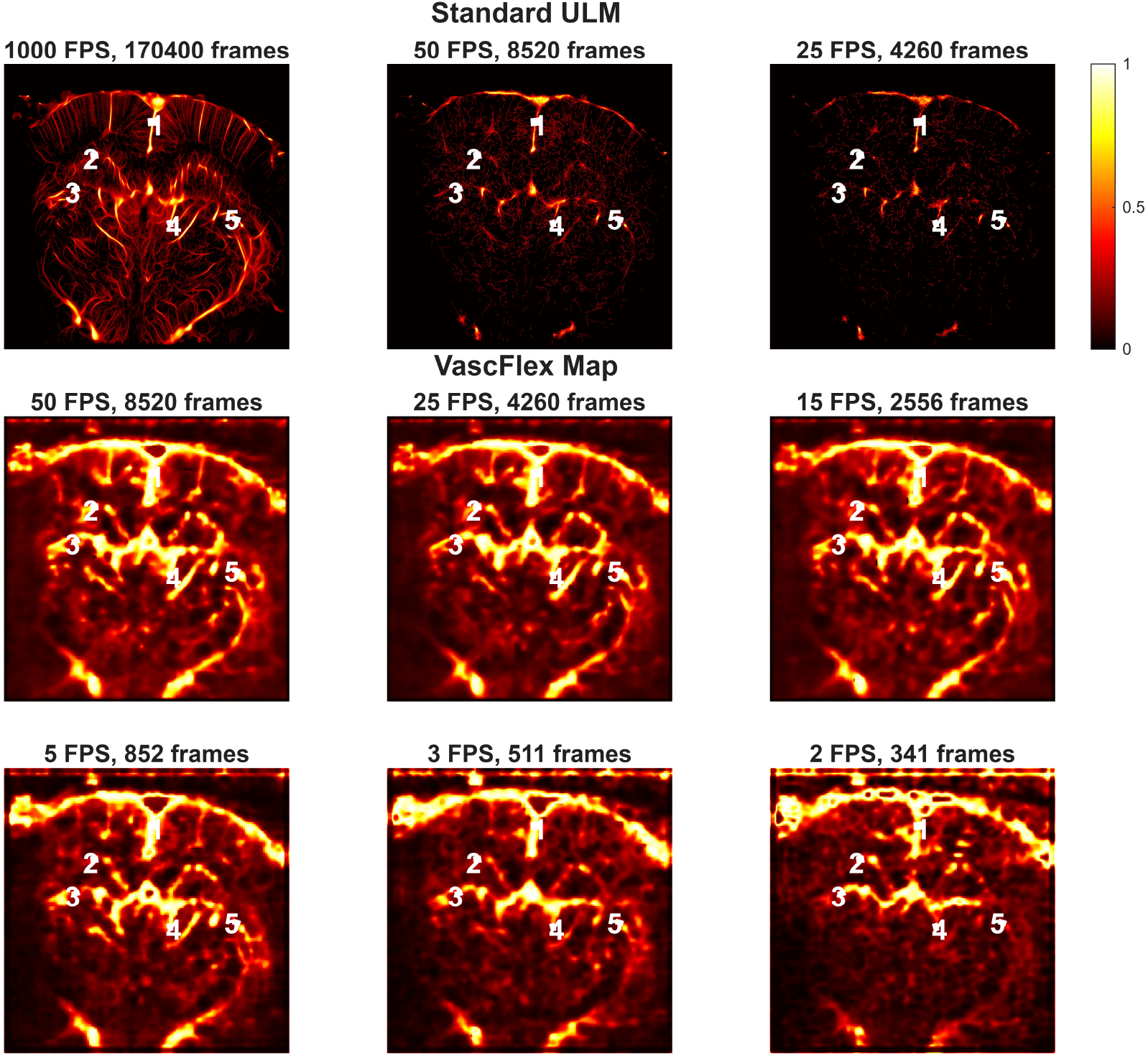
Frame rate ablation study. Row 1: Conventional ULM (1000 FPS). Rows 2–3: VascFlex Map at 50/25/15 FPS and 5/3/2 FPS. White lines indicate FWHM measurement locations (5 vessels).

Fig. 5 examines temporal sufficiency via last-*N* frames analysis. At 25 FPS, vascular recovery improves systematically with longer sequences (2200→200 frames), while 5 FPS maintains topology across 800→200 frames. Both confirm ~5–55 frame minimums suffice for informative vascular mapping.

**Figure 5.**
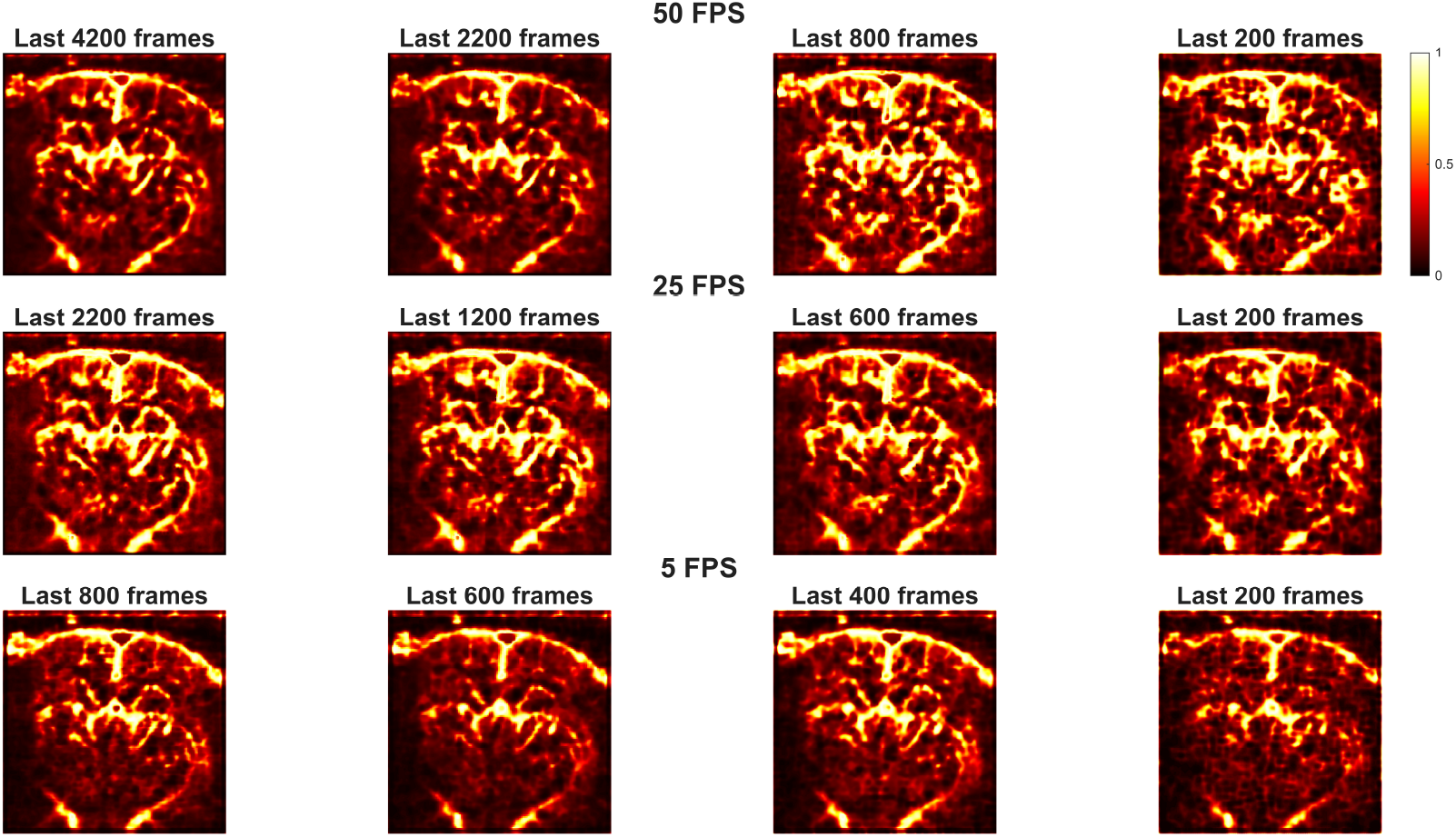
Last-*N* frames analysis demonstrating temporal sufficiency. Both 25 FPS and 5 FPS recover vascular topology from ~200–2200 final frames.

Fig. 6 shows representative −3 dB intensity profiles (Line 1) across all frame rates versus the ULM reference, demonstrating systematic vessel broadening while maintaining consistent localization down to 2 FPS.

**Figure 6.**
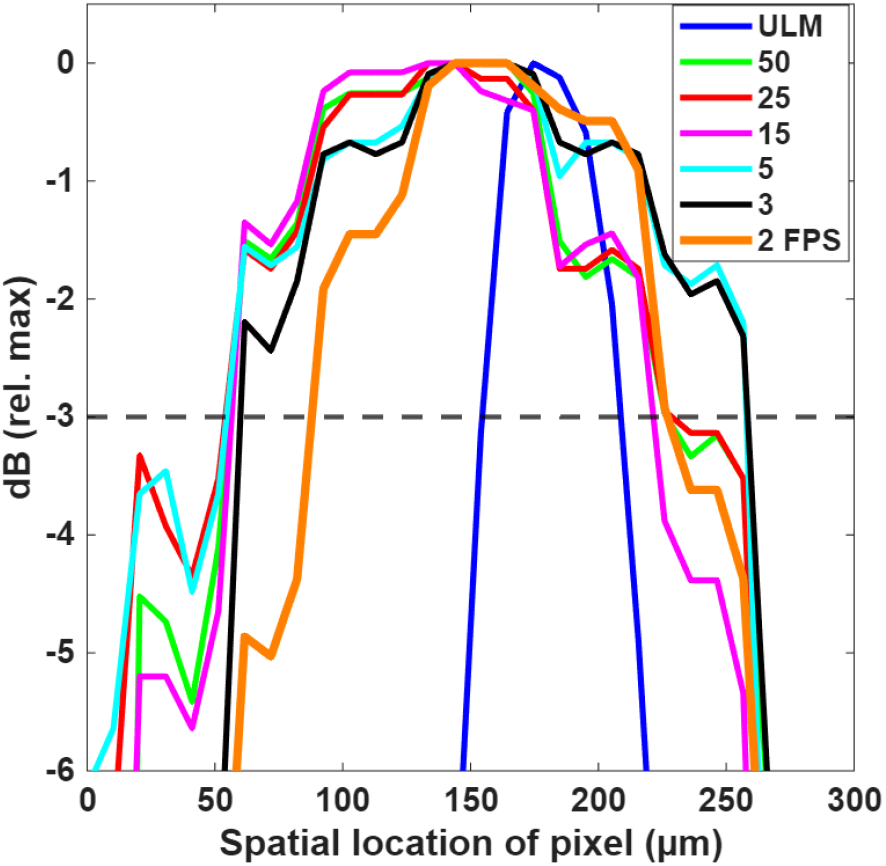
Vessel profile analysis (Line 1). Representative −3 dB profiles across frame rates.

Table 1 summarizes −3 dB FWHM measurements across five vessels (mean ± std). The proposed method achieves 108.83–127.31 *µ*m widths (FPS50–FPS15) versus ULM 34.91 *µ*m (~3–3.6× broadening).

**Table 1:** −3 dB FWHM Across 5 Vessels (*µ*m, Mean ± Std)

| Method | FWHM ($\mu\text{m}$) |
| --- | --- |
| Conventional ULM | $34.91 \pm 30.46$ |
| 50 FPS | $108.83 \pm 43.32$ |
| 25 FPS | $114.99 \pm 35.86$ |
| 15 FPS | $127.31 \pm 24.73$ |
| 5 FPS | $112.93 \pm 47.60$ |
| 3 FPS | $112.93 \pm 50.82$ |
| 2 FPS | $110.88 \pm 15.23$ |

Table 2 presents detailed timing breakdown for the processing pipeline. Fig. 7 shows total end-to-end processing time as a function of the number of frames for different sampling rates. As expected, the total processing time decreases with fewer frames, showing nearly linear scaling for each acquisition rate. For instance, at 25 FPS with 4260 frames, the reconstruction takes approximately 133 s, whereas with 200 frames, the total processing time drops to approximately 28 s, a significant improvement over conventional ULM approaches that typically require hours.

**Table 2:**
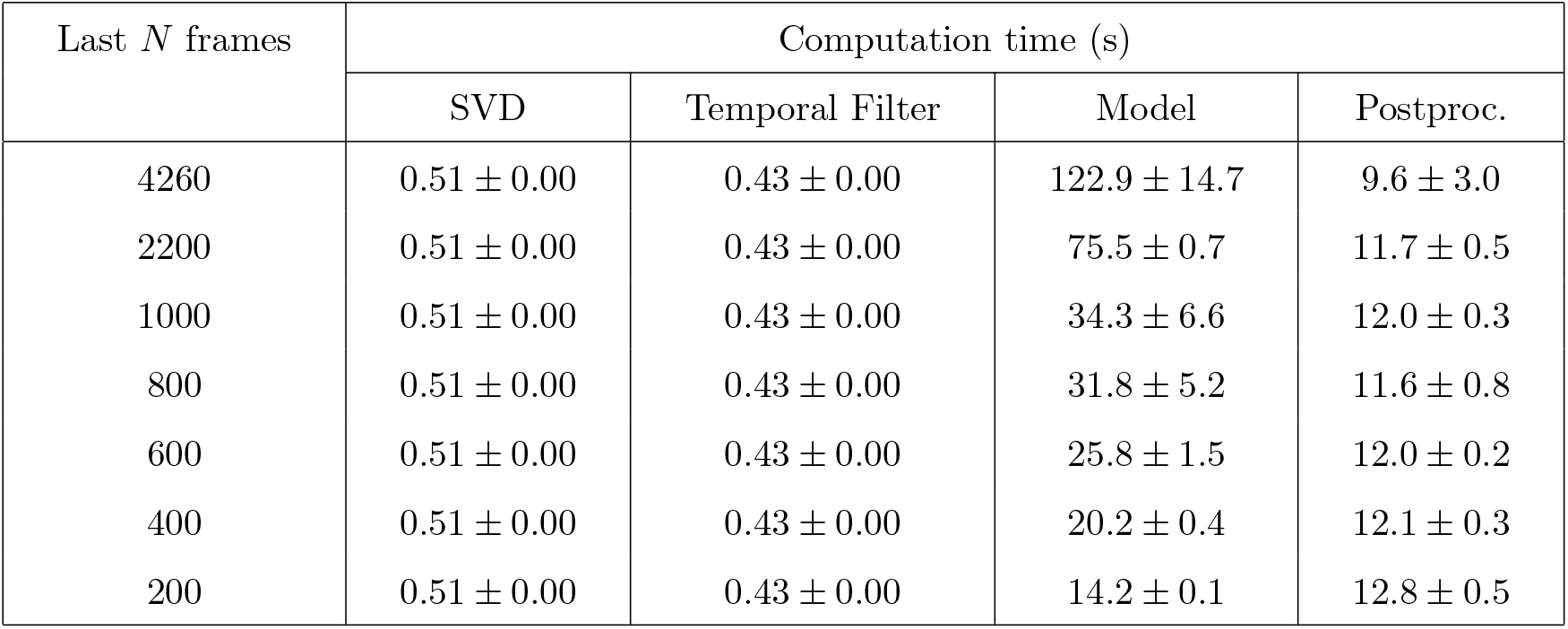
Processing Times (Mean ± Std, Seconds) for 25 FPS Across Last *N* Frames with NVIDIA H100 GPU.

| Last $N$ frames | Computation time (s) | | | |
| --- | --- | --- | --- | --- |
|  | SVD | Temporal Filter | Model | Postproc. |
| 4260 | $0.51 \pm 0.00$ | $0.43 \pm 0.00$ | $122.9 \pm 14.7$ | $9.6 \pm 3.0$ |
| 2200 | $0.51 \pm 0.00$ | $0.43 \pm 0.00$ | $75.5 \pm 0.7$ | $11.7 \pm 0.5$ |
| 1000 | $0.51 \pm 0.00$ | $0.43 \pm 0.00$ | $34.3 \pm 6.6$ | $12.0 \pm 0.3$ |
| 800 | $0.51 \pm 0.00$ | $0.43 \pm 0.00$ | $31.8 \pm 5.2$ | $11.6 \pm 0.8$ |
| 600 | $0.51 \pm 0.00$ | $0.43 \pm 0.00$ | $25.8 \pm 1.5$ | $12.0 \pm 0.2$ |
| 400 | $0.51 \pm 0.00$ | $0.43 \pm 0.00$ | $20.2 \pm 0.4$ | $12.1 \pm 0.3$ |
| 200 | $0.51 \pm 0.00$ | $0.43 \pm 0.00$ | $14.2 \pm 0.1$ | $12.8 \pm 0.5$ |

**Figure 7.**
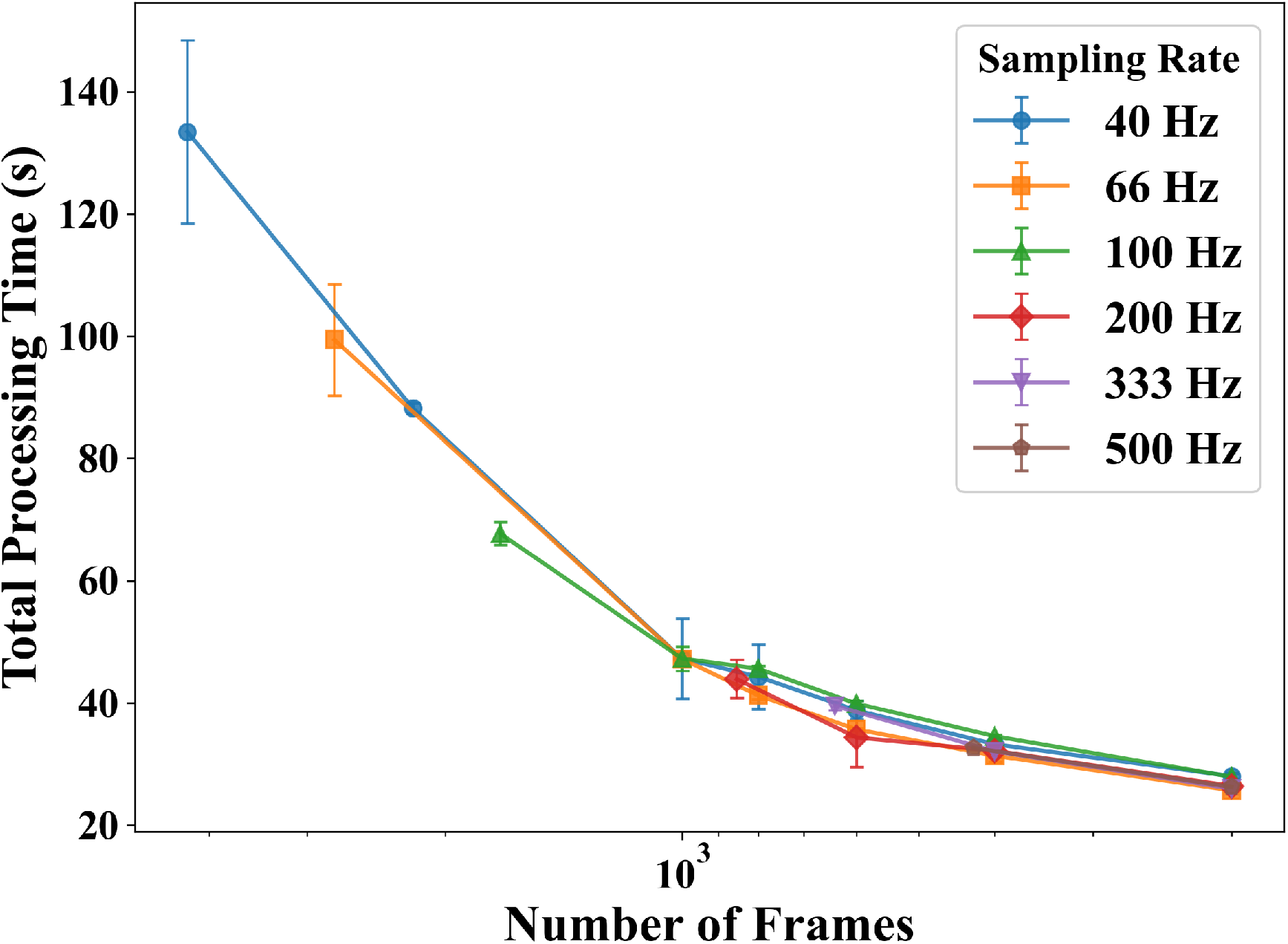
End-to-end computation time scales linearly with input frames. The x-axis is arranged in descending order (high to low frame count, left to right). Processing time decreases from ~133 s (4260 frames) to ~28 s (200 frames), confirming sub-minute reconstruction feasibility across all tested sampling rates.

## 4 Discussion

This work demonstrates that super-resolution-inspired microvascular ultrasound imaging is achievable from data with high temporal sparsity (down to 2 FPS, representing up to 500× data reduction from conventional ULM’s 1000 FPS) using a transformer-decoder architecture with learned linear projections. By operating on low-frame-rate contrast-enhanced ultrasound sequences, VascFlex Map, combined with a post-processing enhancement pipeline, produces coherent vascular maps that preserve clinically meaningful topology without explicit microbubble localization or tracking. Unlike traditional ultrasound localization microscopy (ULM), which is fundamentally dependent on dense temporal sampling for reliable bubble detection, association, and trajectory reconstruction [21, 24], the proposed framework directly learns the spatiotemporal vascular structure from sparse sequences, enabling microvascular visualization under acquisition conditions relevant to standard clinical scanners.

Quantitative analysis confirms a consistent resolution trade-off that reflects the physical limits of sparse sampling. The observed 3–3.6× FWHM broadening relative to ULM (34.91 *µ*m to 108.83– 127.31 *µ*m) represents a predictable loss of the finest capillary-scale detail, yet the reconstructed maps retain higher-order microvascular networks. Importantly, the low variance at extreme sparsity (2 FPS: 110.88 ± 15.23 *µ*m) indicates stable structural recovery rather than stochastic reconstruction, suggesting that the network learns robust vascular priors that generalize across acquisition regimes. This stability is further supported by consistent performance across frame-rate ablation (Fig. 4) and last-*N* frame analyses (Fig. 5), demonstrating that clinically informative vascular topology can be recovered from short sequences and late-bolus frames alone.

The transformer-based temporal encoder plays a central role in enabling this robustness. Global self-attention allows the model to integrate long-range spatiotemporal dependencies across sparse frames, overcoming the limitations of purely convolutional architectures that rely on dense temporal context and local receptive fields [27, 39]. The learned linear projections compress high-dimensional input (262,144 dimensions) to a compact 128-dimensional representation, enabling efficient temporal modeling while the transposed convolutional decoder reconstructs spatial detail. This architecture enables structural coherence even when microbubble observations are temporally discontinuous, explaining the model’s ability to disentangle vascular topology from sparse contrast dynamics and residual clutter.

Computationally, the framework achieves linear scaling with input length and sub-minute inference times across all sparsity regimes (28–133 s on an NVIDIA H100 GPU for 200–4260 frames), representing several orders-of-magnitude acceleration relative to conventional ULM pipelines that require hours of offline processing and multi-gigabyte data handling. Combined with 95%+ data reduction, this approach circumvents one of the primary translational barriers of SR-US: ultrafast hardware dependence, storage and bandwidth constraints, and prohibitive reconstruction latency. The ability to operate directly on low-frame-rate CEUS sequences offers the potential to make the approach compatible with existing clinical platforms.

From the perspective of most clinical applications, the objective of sparse-data microvascular imaging may not be micron-scale capillary resolution, but rapid, interpretable visualization of vascular connectivity, topology, and overall perfusion. The preserved ~100–150 *µ*m-scale vessel architecture may provide sufficient information for detecting ischemia, lesion localization, and vascular pattern recognition in time-critical settings. This approach could also enable a hierarchical imaging paradigm: rapid low-frame-rate sparse reconstruction for global perfusion mapping and region-of-interest identification, followed by targeted high-fidelity ULM acquisition where ultra-high resolution is clinically justified.

## 5 Study Limitations

Several limitations of the proposed framework are acknowledged. The reconstructed vessel maps exhibit approximately 3–3.6× FWHM broadening relative to conventional ULM, producing vascular topology maps rather than true super-resolution capillary detail. The method generates static vessel probability maps and does not provide blood flow velocity or directionality information available from conventional ULM through microbubble tracking. The current evaluation employs motion-free datasets, and robustness to physiological motion artifacts such as respiration and cardiac pulsation remains to be assessed in future work.

Despite these limitations, VascFlex Map enables rapid vessel architecture mapping and could be compatible with standard clinical ultrasound systems. This approach could find applications in stroke triage, tumor margin assessment and classification, and vascular screening, where dense ultrafast acquisition is impractical. This approach could also serve as a “first cut” reconstruction for SR-US, to ensure data fidelity, and subsequent high-resultion reconstructions could be performed offline. Future work will focus on resolution enhancement, volumetric 3D extensions for whole-organ mapping, integration of quantitative biomarkers and real-time deployment on ultrasound imaging platforms.

## 6 Conclusion

This study introduced **VascFlex Map**, a transformer-decoder framework with learned linear projections that enables practical microvascular ultrasound imaging from low-frame-rate, sparsely sampled CEUS data. By reconstructing vascular probability maps directly from 2–50 FPS sequences, the method, followed by a post-processing enhancement stage, achieved coherent microvascular recovery under 20–500× data reduction relative to conventional ULM. Evaluation on *in vivo* rat brain bolus data demonstrated preserved vascular maps down to 2 FPS (341 frames), with −3 dB FWHM of 110.88±15.23 *µ*m compared to 34.91±30.46 *µ*m for full-data ULM, reflecting an expected resolution trade-off while maintaining key details.

The framework delivered sub-minute reconstruction times (28–133 s on NVIDIA H100 GPUs), eliminating dependence on ultrafast acquisition hardware, and reduced data storage and transfer requirements by over 95%. The convergence of computation times across higher sampling rates for smaller frame counts indicates efficient parallelization on the GPU. Overall, these findings confirm that sub-minute per-frame reconstruction is achievable even with large datasets. These characteristics directly address the three principal translational barriers of super-resolution ultrasound: specialized hardware requirements, massive data volumes, and prohibitive computational latency.

## Acknowledgment

The authors thank the PALA challenge organizers for providing the rat brain bolus dataset. This work was supported by IIT Gandhinagar and the Ministry of Education STARS (Grant No. MoE-STARS/STARS-2/2023-0620).

